# Multimodal interrogation of ventral pallidum projections reveals projection-specific signatures and opposite roles in cocaine withdrawal

**DOI:** 10.1101/2021.11.15.468637

**Authors:** Nimrod Bernat, Rianne Campbell, Hyungwoo Nam, Mahashweta Basu, Tal Odesser, Gal Elyasaf, Michel Engeln, Ramesh Chandra, Shana Golden, Seth Ament, Mary Kay Lobo, Yonatan M. Kupchik

## Abstract

The ventral pallidum (VP) is central to reward seeking and withdrawal from drugs of abuse. A characteristic of the VP is the diversity of its projection targets. Yet, it remains unknown whether different VP projections also differ in other aspects, such as their transcriptome, physiology and relevance to drug reward. In this study we perform a multimodal dissection of four major projections of the VP – to the lateral hypothalamus (VP_→LH_), ventral tegmental area (VP_→VTA_), lateral habenula (VP_→LHb_) and mediodorsal thalamus (VP_→MDT_) – with physiological, anatomical, genetic and behavioral tools and show significant differences between projections in all aspects. Specifically, the VP_→LH_ and VP_→VTA_ projections show minimal overlap and stand out as having opposite properties – VP_→LH_ neurons show higher excitability compared to VP_→VTA_ neurons, different pattern of inputs and differentially expressed genes. Moreover, inhibition of VP_→LH_ projections diminishes, while inhibition of VP_→VTA_ enhances cocaine preference after cocaine withdrawal. This demonstrates that VP projections are heterogenous neuron populations with different roles in cocaine withdrawal.

## Introduction

The ventral pallidum (VP) is a central hub in the basal ganglia and plays important roles in motivation, reward, and addiction^1–3^. It consists mainly of GABAergic neurons, with a minority of glutamatergic and cholinergic neurons^1^. Its neurons are also sometimes categorized by a neuropeptide they release^4^ or the expression of a particular receptor^5^ or protein^1,2,6,7^.

In addition to the intricate cellular composition, the VP also shows complex connectivity. It receives mainly inhibitory inputs from various limbic regions^1^ while having a wide range of downstream targets, including dense projections to the ventral tegmental area (VP_→VTA_), the lateral hypothalamus (VP_→LH_), the mediodorsal thalamus (VP_→MDT_) and the lateral habenula (VP_→LHb_)^1^. Previous work has shown that VP projection neurons exert different roles in rewarding and aversive behaviors^1,8–12^. Despite these distinct roles in behavior, it is theorized that different VP projections are composed largely of the same neuronal composition, possibly even with significant overlap between the projection subpopulations.

The development of new technologies such as rabies-based tracing^13,14^, single-cell RNA sequencing^15^ and cell-specific optogenetics^16–18^ or chemogenetics^19^ reveals in recent studies an increasing heterogeneity of neurons residing within the same brain region, defined by their connectivity patterns, gene expression, physiology, morphology and more^20,21^. The high complexity of these methods has driven most studies to focus on one modality when analyzing the heterogeneity of neurons in a certain region. In particular, heterogeneity is often described either based on intracellular properties (gene expression, physiology, etc.) or on system-level connectivity, without integrating these two levels of complexity. As neurons are multimodal entities there is a need for a multimodal interrogation of neuronal heterogeneity.

In this collaborative work we integrate multilevel approaches to examine whether the different projections of the VP to the VTA, LH, MDT and LHb are separate neuronal subpopulations based on their inputs, physiology, transcriptome and behavioral relevance. Our data highlight the VP_→VTA_ and VP_→LH_ projections as highly distinct in these parameters, suggesting these two projections of the VP may be different types of neurons.

## Methods

### Experimental subjects

For electrophysiology, neural tracing and behavioral experiments, subjects were naïve C57bl6/J wildtype male and female mice who were approximately 8 weeks old at the beginning of the experiments. A 12-hour reversed light/dark cycle was maintained, with the lights turned off at 8:00 am. Experimental procedures were conducted during dark hours. Mice were group-housed and nesting/enrichment material was available. Experimentation began after a minimum of 7 days of acclimation to animal facility. All experimental procedures were approved by the Authority for Biological and Biomedical Models in the Hebrew university. For projection-specific gene expression analysis, adult (7-8 weeks old) male RiboTag (RT)+/+ mice (Rpl22tm1.1Psam/J) on a C57Bl/6 background were used. RiboTag mice were given food and water ad libitum throughout the study. These studies were conducted in accordance with guidelines of the Institutional Animal Care and Use Committees at University of Maryland School of Medicine.

### Stereotaxic microinjections

Mice were anesthetized with isoflurane and fixed in a stereotaxic frame (Kopf, Model 940). Viruses or red Retrobeads^™^ (Lumafluor, Durham, North Carolina) were microinjected into one or more of the following structures - the MDT (coordinates in millimeters relative to Bregma: anterior/posterior −1.5, medial/lateral ±0.47, dorsal/ventral −3.63), LHb (A/P −1.8, M/L ±0.7, D/V −2.91), LH (A/P −0.3, M/L ±0.9, D/V - 5.5), VTA (A/P −2.9, M/L ±0.5, D/V −4.53), VP (A/P 0.7, M/L ±1.2, D/V −5.21) and NAc (A/P 1.8, M/L ±1.0, D/V −4.6) (see below for details on the injections in each experiment). Injections were performed bilaterally by drilling bilateral holes into the skull and then microinjecting the viral constructs [through a 33 ga NanoFil syringe (World Precision Instruments; 300 nl per hemisphere, 100 nl/min, needle retracted 5 min after injection terminated)] or Retrobeads [through a 30 ga syringe (Hamilton: 300 nl per hemisphere, 300 nl/min, needle retracted 5 min after injection terminated)] into the target structure.

Circuit-specific gene expression analysis was performed using retrograde AAV-Cre virus in combination with Ribotag mice. Briefly, anesthetized RiboTag RT +/+ mice were bilaterally injected with AAV5-Cre viruses (AAV sterotype 5 AAV5.hSyn.HI.eGFP-Cre.WPRE.SV40 (Addgene; #105540) at the following coordinates: VP (from Bregma; 10° angle, anterior/posterior: AP +0.9, medial/lateral: ML ±2.2, dorsal/ventral: DV −5.3), MDT (from Bregma; 10° angle, AP: −0.8, ML: ±1.2, DV: −3.7), VTA (from Bregma; 7° angle, AP: - 3.2, ML: ±1, DV: −4.6), LHb (from Bregma; 10° angle, AP: −1.2, ML: ±0.7, DV: - 3.1) and LH (from Bregma; 10° angle, AP: −0.7, ML: ±1.4, DV: - 5.2).

### Double-labeling experiments

Each mouse was injected with two different retrograde tracers, rAAV-GFP and rAAV-RFP (ELSC viral core, The Hebrew University, Israel). Each virus was injected bilaterally (300 nl per hemisphere) into one of the VP targets tested here. Two weeks after microinjections mice were anesthetized with Pental (CTS Chemical Industries, Israel) and perfused with 4% paraformaldehyde (PFA). Sections of the VP (40 μm) were prepared using a sliding microtome (Leica, model SM2010R) and mounted on slides. The sections were scanned with Nikon ECLIPSE-NiE fluorescent microscope and scans were analyzed using ImageJ (NIH).

### Whole-brain mapping of inputs to VP projections

To label monosynaptic inputs to specific VP projections we used the “<u>T</u>he <u>R</u>elationship between <u>I</u>nputs and <u>O</u>utputs” (TRIO) technique^13^. Mice were first injected with a retrograde virus expressing Cre recombinase (rAAV-Cre; ELSC viral core, The Hebrew University, Israel) into one of the four targets of the VP examined here (300 nl). We also injected these mice with a cocktail of two helper viruses expressing the TVA receptor (AAV-FLEx-TVA-mCherry) and the rabies glycoprotein (AAV-FLEx-RG) (total of 500 nl, 250 nl per virus) into the VP. Two weeks after these injections we injected mice with the modified rabies virus (500 nl) into the VP. We allowed the virus 7 days to spread before anesthetizing and perfusing the mice with PFA as described above. Brains were then post-fixed with 4% PFA overnight at 4 °C. Tissue clearing and staining were performed according to the detailed protocol available at http://idisco.info. Briefly, brains were dehydrated, bleached to reduce background autofluorescence, rehydrated, then incubated with rabbit anti-RFP antibody (1:1000, Rockland, Cat# 600-401-379) and chicken anti-GFP antibody (1:2000, Aves labs, Ca# GFP1010) for 5 days, followed by washing and incubation in Cy3 conjugated anti rabbit secondary antibody (1:500, Jackson ImmunoResearch, Cat# 711-165-152) and Cy5 conjugated anti chicken secondary antibody (1:400, Jackson ImmunoResearch, Cat# 703-175-155) for 5 days. Following additional washing, brains were dehydrated and cleared in methanol and dichloromethane, then refractive index matching occurred in dibenzyl ether. Brains were imaged at x4 magnification, 0.35 numerical aperture, in the vertical orientation on a LaVision Ultramicroscope II lightsheet microscope with a near-isotropic xyz resolution of 1.6 μm × 1.6 μm × 2 μm. Images were acquired with a 488 nm laser for an autofluorescence reference channel, and 561 nm and 642 nm lasers for acquisition of secondary antibody fluorescence. The ClearMap Python package (www.github.com/christophkirst/clearmap) was used for cell detection and registration of cell coordinates onto the Allen Brain Atlas. Data analysis occurred on an Antec P110 Silent Workstation running Ubuntu 20.04LTS, and followed steps outlined in ClearMap documentation. Briefly, data were downsampled and the Elastix toolbox (http://elastix.isi.uu.nl/) was used to perform automated 3D affine and B-spline transformation to register the autofluorescence signal channel to the 25 μm resolution Allen Brain Atlas reference, and to correct for any motion between imaging of the autofluorescence and signal channel images. Elastix’s ‘pixel classification’ machine learning workflow was used to teach a model to segment input neurons from background, which later was used in Headless mode in ClearMap to detect input neurons. The Transformix module of the Elastix toolbox was then used to apply transformation vectors from the registration step to cellular coordinates, and cell counts for each region were calculated.

### RNA-Sequencing and Bioinformatics

Three weeks following viral injections, VP tissue was collected from Ribotag mice infused with Retrograde-Cre virus (AAV5-Cre). Projection-specific RNA isolation was performed using polyribosome immunoprecipitation as described previously^22,23^. Briefly, pooled tissue from RiboTag (RT)+/+ mice with virally mediated Cre expression in VP projection neurons (n= 5 mice per sample) or in VP was homogenized and 800 μL of the supernatant was incubated in HA-coupled magnetic beads (Invitrogen: 100.03D; Covance: MMS-101R) overnight at 4°C. Magnetic beads were washed using high salt buffer. Following TRK lysis buffer, RNA was extracted with the RNeasy Micro kit (Qiagen: 74004). Libraries were prepared from 10 ng of RNA using the NEBNext Ultra kit (New England BioLabs, Ipswich, MA, USA) and sequenced on an Illumina HiSeq 4000 with a 75 bp paired-end read. RNA-sequencing data are available through the Gene Expression Omnibus database (GEO accession number: GSE218580).

A total of 75–110 million reads were obtained for each sample. Reads were aligned to the mouse genome using TopHat and the number of reads that aligned to the predicted coding regions was determined using HTSeq. Significant differential expression was assessed using Limma by comparing VP projection neuron samples to total VP. Genes with the absolute value of log fold change (LFC)≥0.3 and an uncorrected p-value<0.05 in the pairwise comparisons were used for downstream analysis (**Supplementary Table 1**). Cytoscape 3.7.2 software was used for downstream analysis: transcriptional regulator networks were identified using the iRegulon app and Gene Ontology functional enrichment analysis was performed using the BiNGO app (**Supplementary Table 2**). From these lists, top GO terms were selected based off of highest −log_10_(adjusted-p-value), and top upstream regulators were selected based on the number of predicted targets DEGs.

### Slice preparation

As described before ^24–26^. Mice were anesthetized with 150 mg/kg Ketamine-HCl and then decapitated. Coronal slices (200 μm) of the VP were prepared (VT1200S Leica vibratome) and moved to vials containing artificial cerebrospinal fluid (aCSF (in mM): 126 NaCl, 1.4 NaH_2_PO_4_, 25 NaHCO_3_, 11 glucose, 1.2 MgCl_2_, 2.4 CaCl_2_, 2.5 KCl, 2.0 Na Pyruvate, 0.4 ascorbic acid, bubbled with 95% O_2_ and 5% CO_2_) and a mixture of 5 mM kynurenic acid and 100 mM MK-801. Slices were stored in room temperature (22-24 °C) until recording.

### In vitro whole-cell recording

Recordings were performed at 32 °C (TC-344B, Warner Instrument Corporation). VP neurons were visualized using an Olympus BX51WI microscope and recorded from using glass pipettes (1.3-2.5 MΩ, World Precision Instruments) filled with internal solution (in mM: 68 KCl, 65 D-gluconic acid potassium salt, 7.5 HEPES potassium, 1 EGTA, 1.25 MgCl_2_, 10 NaCl, 2.0 MgATP, and 0.4 NaGTP, pH 7.2-7.3, 275mOsm). Multiclamp 700B (Axon Instruments, Union City, CA) was used to record both membrane and action potentials and postsynaptic currents (PSCs) in whole-cell configuration. Excitability and passive membrane properties were measured in current clamp mode while synaptic activity was measured in voltage clamp mode at a holding potential of −80 mV. Recordings were acquired at 10 kHz and filtered at 2 kHz using AxoGraph X software (AxoGraph Scientific, Sydney). To evoke inhibitory PSCs (IPSCs) from NAc terminals expressing ChR2 we used a 470 nm LED light source (Mightex Systems; 0.1–1 ms in duration) directed at the slice through the x60 objective. The stimulation intensity was set to evoke 50% of maximal IPSC at −80 mV. Recordings were collected every 10 seconds. Series resistance (Rs) measured with a −2 mV hyperpolarizing step (10 ms) given with each stimulus and holding current were always monitored online. Recordings with unstable Rs or when Rs exceeded 20 MΩ were aborted.

### Current clamp experiments

After penetrating the neuron, we switched to current clamp mode and recorded the resting membrane potential of the neurons and spontaneous action potentials for 60 seconds. Cells with unstable membrane potential were discarded. Action potentials were later detected by their waveform using the Axograph software, baselined and analyzed. Potentials of <20 mV in amplitude were discarded. We then applied the current step protocol - five 500 ms-long depolarization current steps ranging from 0 pA to +80 pA (20 pA intervals) were applied, inter-step interval was 3 seconds. The 5-step protocol was repeated 5 times with 3 s between repetitions. Baseline membrane potential was adjusted to be approximately −50 mV in all neurons by injecting constant current.

### Cocaine conditioned place preference

Mice were first injected with rAAV-Cre in one of the 4 VP targets examined here and with an AAV encoding for the Gi-coupled DREADD, hM4Di, in a Cre-dependent manner (AAV-DIO-hM4Di-mCherry), or a sham virus (AAV2-EF1a-DIO-EYFP-WPRE-pA) into the VP. After two weeks of acclimation to the reverse light cycle and recovery from surgery, mice were trained in the unbiased cocaine conditioned place preference (CPP) paradigm as described previously ^24^ (**Fig. 8**). On the first day, mice were allowed to freely explore both sides of a 30 cm × 30 cm arena, divided in two, each side with a different wall pattern and floor texture. On the following days, experimental mice received one daily injection of either cocaine (15 mg/kg, i.p.) or saline such that cocaine was always given in one side of the box (the “cocaine-paired” side) and saline in the other. Cocaine-paired sides were counterbalanced for pattern and side. Cocaine/saline injections alternated daily until each mouse received 4 of each. Control mice received 8 injections of saline. Then, mice were left in their cages for 14 days (abstinence) before being tested for their preference of the cocaine-paired side. On the test day, mice first received an injection of the DREADD ligand clozapine-N-oxide (CNO, 3 mg/kg, i.p.), were left in their home cages for 30 minutes and then put in the CPP box for 15 minutes. During the test, mice were allowed to move freely between the two sides of the arena. Movement was tracked using MediaRecorder (Noldus, the Netherlands), analyzed using Optimouse software^27^ and CPP scores were calculated offline as the ratio between the difference in time spent between the cocaine-paired and unpaired sides and the total time [CPP score = (time in paired zone - time in unpaired zone)/(time in paired zone + time in unpaired zone)].

### Viruses

Viruses –
| Viral construct | manufacturer | Abbreviation in paper | titer | Used in figure # |
| --- | --- | --- | --- | --- |
| AAV5.hSyn.HI.eGFP-Cre.WPRE.SV40 | Addgene | AAV5-Cre | 7.0E+12 | 2,3,7 |
| AAVr-hSyn-eGFP | EVCF | rAAV-GFP | 1.78E+13 | 1 |
| AAVr-hSyn-mRuby2 | EVCF | rAAV-RFP | 3.06E+13 | 1 |
| AAVr-hSyn-iCre | EVCF | rAAV-Cre | 1.10E+14 | 4,8 |
| EnvA G-Deleted Rabies-eGFP | Salk inst. | GdRV-GFP | 2.91E+07 | 4 |
| AAV-CA-FLEX-RG | UNC | AAV-FLEX-RG | 3.2E+12 | 4 |
| AAV-EF1a-FLEX-TVA-mCherry | UNC | AAV-FLEX-TVA | 4.3E+12 | 4 |
| AAV2-EF1a-DIO-hChR2(H134R)-EYFP | UNC | AAV-DIO-ChR | 5.7E+12 | 5 |
| AAV2-hSyn-DIO-hM4D(Gi)-mCherry | UNC | AAV-DIO-DREADDs | 3.7E+12 | 8 |
| EVCF – ELSC Vector Core Facility, The Hebrew University of Jerusalem; Salk – Salk Institute Viral Vector Core; UNC – University of North Carolina Vector Core |  |  |  |  |

### Statistics and data analysis

All statistical analyses were performed using GraphPad Prism 9.2 (GraphPad Software Inc.). Parametric tests were used unless stated otherwise with p values <0.05 considered as significant. All tests are indicated in the figure legends. Data in graphs represent mean ± s.e.m.

## Results

### Minimal overlap between VP projection neurons

While the VP projects to the LH, VTA, MDT and LHb, it is not known whether there is any overlap between these projections, and if so to what extent. To test this we performed six experiments examining the overlap between all possible pairs of the four targets of the VP. For each pair combination we injected one retrograde virus expressing GFP (rAAV-GFP) into one of the targets and another retrograde virus expressing RFP (rAAV-RFP) into the other target (**Fig. 1A**). Then we examined the proportion of the double-labeled yellow neurons out of all labeled neurons in the VP (**Fig. 1B**). Our data show that overlap between projections ranged between 7-20%, with the VP_→LH_ projections showing the lowest overlap proportions and especially low overlap with VP_→VTA_ projections (7%) (**Fig. 1C**). This may be surprising, as a previous study^28^ showed that most of the VP projection neurons, regardless of their final target, pass through the LH, but another study showed different behavioral effects for VP_→LH_ and VP_→VTA_ projections^11^. It is possible that the levels of overlap between VP projections reported here are underestimations as the overlap may be affected by the efficiency of virus uptake by axons and by microinjection localization.

**Figure 1.**
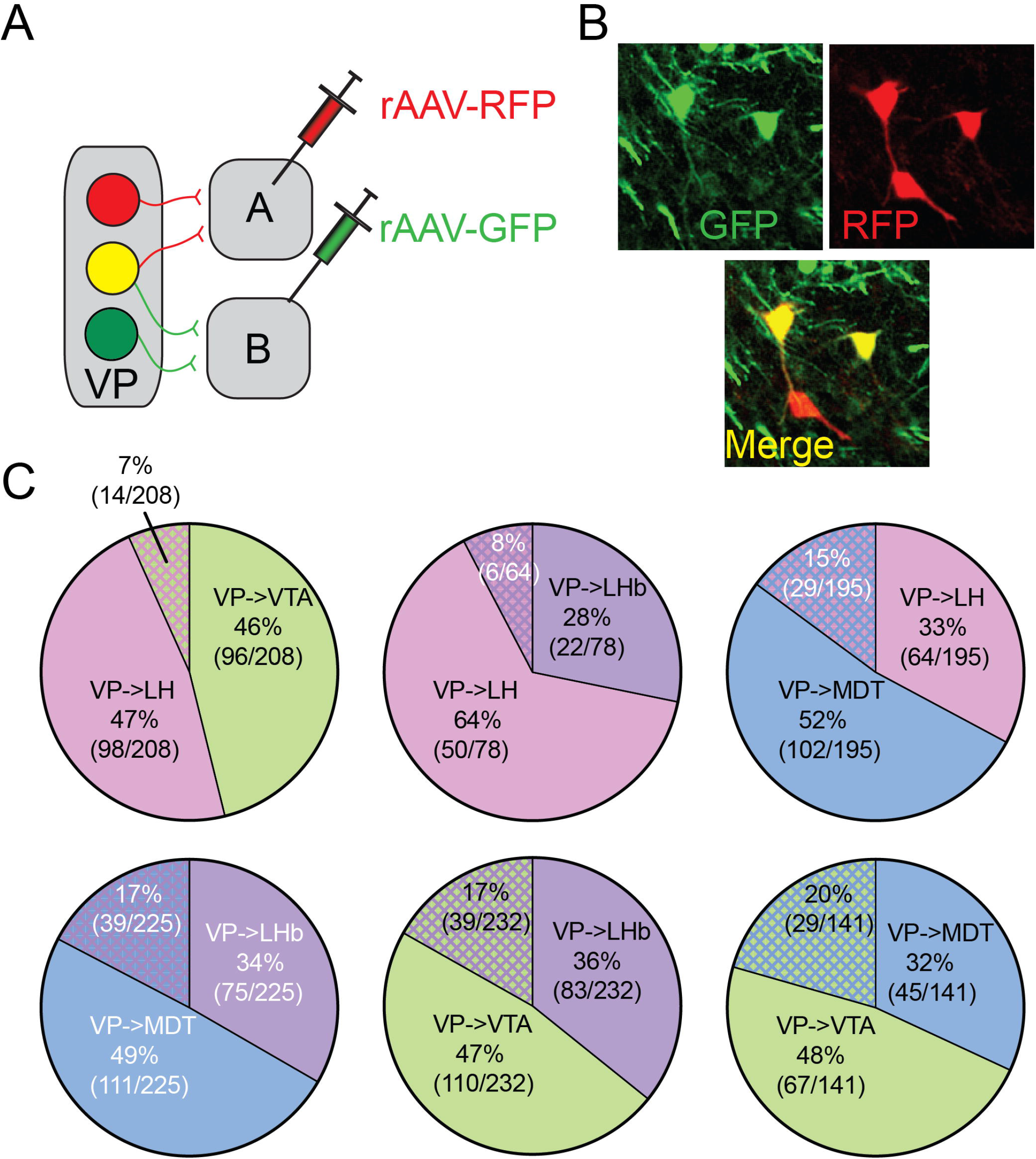
Minimal overlap between ventral pallidal projections to the MDT, LHb, LH and VTA. (**A**) Experimental setup. We injected each mouse with two retrograde viruses expressing either RFP (rAAV-RFP) or GFP (rAAV-GFP), each in a different target of the VP. Thus, we could identify neurons projecting to either target and those projecting to both. (**B**) Sample images of labeled VP_→MDT_ neurons, VP_→LH_ neurons and double-labeled neurons projecting to both targets. (**C**) Pie charts depicting for each pair of VP targets the proportion of double-labeled VP neurons projecting to both targets out of all labeled neurons (arranged from lowest to highest proportion of double-labeled neurons). The proportion of VP neurons projecting to both targets tested was in the range of 7-20%. Note that VP_→LH_ neurons showed the lowest proportions of overlap, especially with VP_→VTA_ neurons.

### Distinct molecular signatures within VP projection neurons

Given that the projections of the VP originate in largely distinct sets of neurons, we aimed to examine whether these sets of VP projection neurons exhibit distinct profiles at the molecular level. Although studies have identified the neurochemical composition of VP projection neurons using *in situ* hybridization^8,9,29^, we profiled global translatome-wide signatures within VP projection neurons. This allows us to identify patterns and novel genes enriched within VP projection neurons.To this end, gene expression profiles of VP projection neurons were assessed using RNA-sequencing (**Fig. 2A**). Retrograde Cre virus (AAV5-Cre) was infused either into the VTA, LHb, LH, MDT or VP. This resulted in Cre-dependent expression of HA-tagged ribosomal protein within upstream VP neurons. Infusion of AAV5-Cre directly into the VP served as a comparison of non-pathway specific VP (global VP) gene expression. VP tissue was collected from mice and Ribotag methods were used to isolate pathway-specific ribosome-associated mRNA from VP neurons. RNA-sequencing libraries were constructed from each VP projection group and followed by translatome profiling (**Fig. 2A**).

**Figure 2.**
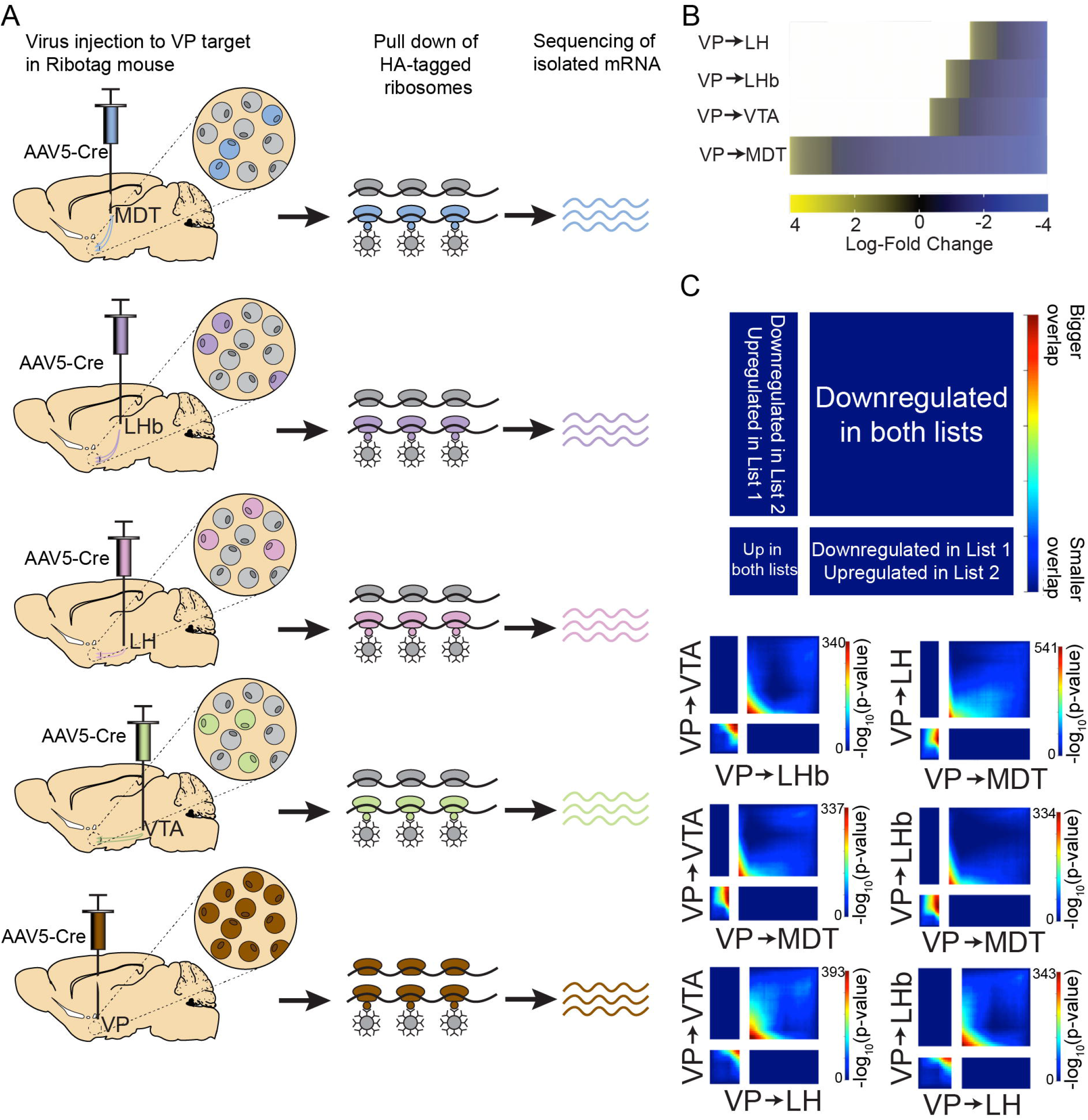
VP projection neurons have distinct molecular signatures. (**A**) Retrograde Cre (AAV5-Cre) was infused into 1 of 4 downstream VP regions (MDT; LHb;LH;VTA) or VP within Ribotag male mice (n=16 mice per projection). HA-immunoprecipitation procedures were employed to isolate RNA from distinct VP projection neurons or all VP neurons. RNA-sequencing libraries were generated and analyzed from these samples to characterize baseline gene expression profiles within VP neurons projecting to the MDT, LHb, LH, VTA. (**B**) Differential Gene Expression (DEG) analysis was generated by comparing gene expression within specific VP projection neuron types relative to the global VP gene expression (uncorrected p<0.05). Transcriptional patterns within each VP projection neuron type are shown in a heat map. (**C**) Rank-Rank Hypergeometric Overlap (RRHO) analysis comparing gene expression in a threshold free manner suggests VP_→MDT_/VP_→LH_ and VP_→MDT_/VP_→LHb_ projection neurons have the highest concordance of gene expression patterns.

First, pair-wise comparisons of gene expression patterns from VP_→VTA_, VP_→LHb_, VP_→LH_ and VP_→MDT_ were compared to gene expression within non-projection specific VP neurons (**Fig. 2B**). The following number of differentially expressed genes (DEGs) were detected in each VP projection neuron: VP_→MDT_: 4318; VP_→VTA_: 1982; VP_→LH_: 1709 and VP_→LH_: 1308. To further characterize global gene expression patterns, we performed Rank-Rank Hypergeometric Overlap (RRHO) analysis^30^, which compares gene expression between two lists in a threshold-free manner. Heat maps display overlap at those points, determined by relative effect sizes in differential gene expression using - log_10_(p-value). Comparisons between gene expression lists of all six possible pairs of VP projections were performed (**Fig. 2C**). High concordance is seen within genes upregulated within VP_→LH_ versus VP_→MDT_ in relation to VP global neurons (peak: −log_10_(p-value): 541) and VP_→LHb_ versus VP_→MDT_ (peak: −log_10_(p-value): 334). High concordance is also observed within downregulated genes in VP_→VTA_ versus VP_→LH_ neurons (peak: −log_10_(p-value): 393). These data overall suggest that VP_→MDT_ has the highest number of upregulated genes and exhibits the highest overlap of upregulated gene expression patterns among other VP projection neuron types.

To identify cell-type specific molecules and processes, additional analysis was performed on genes upregulated (i.e. enriched) within the VP projections. VP_→MDT_ neurons had the highest number of upregulated DEGs (708) in comparison to other VP neuron types (VP_→VTA_: 489; VP_→LH_: 454; VP_→LHb_: 402). Upregulated genes were largely distinct from one another (**Fig. 3A**), with only 66 upregulated genes shared amongst all VP neuron types. Consistent with the patterns detected from RRHO analysis, VP_→MDT_ and VP_→LH_ projection neurons shared the highest number of upregulated genes (267 total) and VP_→LH_ and VP_→LHb_ have little gene overlap (133 total). To characterize the biological processes and cellular functions associated with genes enriched within each VP neuronal type, gene ontology (GO) analysis was performed (**Fig. 3B**). All projection neuron types have enrichment for GABA receptor activity processes, suggesting that all these neuronal types receive GABAergic inputs. VP_→VTA_ neurons have genes enriched for glutamate receptor activity and VP_→LHb_ neurons are enriched for dopamine binding, which were not detected in other projection neuron types. Interestingly, shared GO terms across VP projection types include synaptic transmission and cytoskeletal protein binding. This may indicate that VP projection neurons express distinct sets of genes to regulate processes for synaptic communication. Similarly, GO terms for metabolic process and mitochondrion organization are detected amongst VP_→VTA_ and VP_→MDT_ neurons. This suggests that neuronal types may have distinct molecules that regulate mitochondrial and energy processes. Overall, the enriched genes and biological processes within each projection neuron type may underlie the VP projection-specific functions and responses.

**Figure 3.**
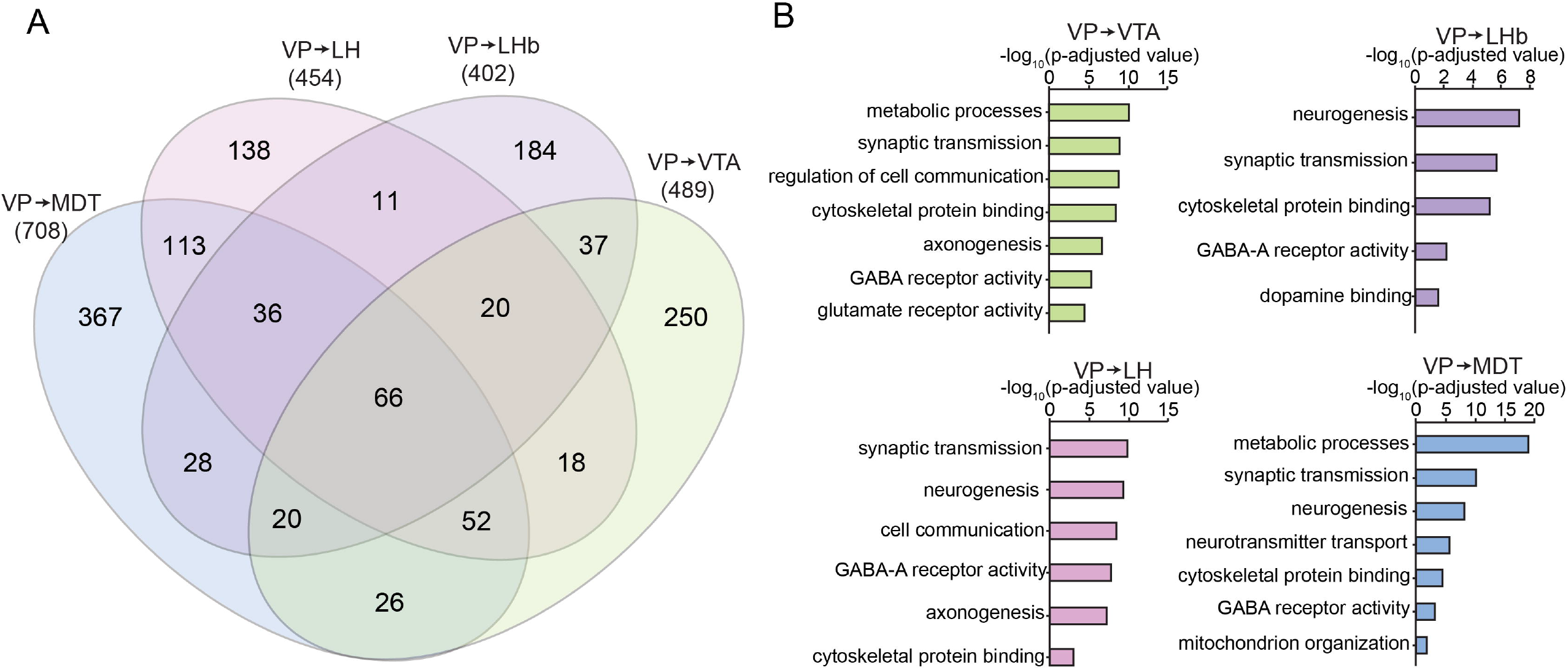
Gene ontology analysis identifies biological processes and molecular functions enriched within VP projection neurons. (**A**) Genes upregulated within each VP projection neuron type were compared in Venn diagrams. The VP_→MDT_ projection neuron has the highest number of enriched genes, with VP_→MDT_ and VP_→LH_ sharing the highest number of enriched genes. (**B**) Top Gene Ontology (GO) terms enriched in upregulated genes from each VP projection neuron.

### Different input patterns for different VP projections

The four VP projections we test here differ, by definition, in the target they project to. However, they may also receive their inputs from different sources, thus generating parallel segregated circuits. To test whether the VP projections differ in their input patterns we used the Testing the Relationship between Input and Output (TRIO) method^13^, which allows labeling of monosynaptic inputs to a specific projection neuron (**Fig. 4A**). We first injected a retrograde virus encoding Cre recombinase (rAAV-Cre) into either of the four VP targets (i.e. VTA, LH, MDT or LHb) and then viruses that encode for the TVA receptor and the rabies glycoprotein in a Cre-dependent manner into the VP. Thus, only VP neurons projecting to the site injected with the rAAV-Cre would express the TVA receptor and the rabies glycoprotein (“starter cells”, **Supplementary Figure 1**). Lastly, we injected the modified rabies virus^31^ into the VP and tested neuron labeling using the iDisco+ method^32^ and light sheet fluorescence microscopy (**Fig. 4A,B**).

**Figure 4.**
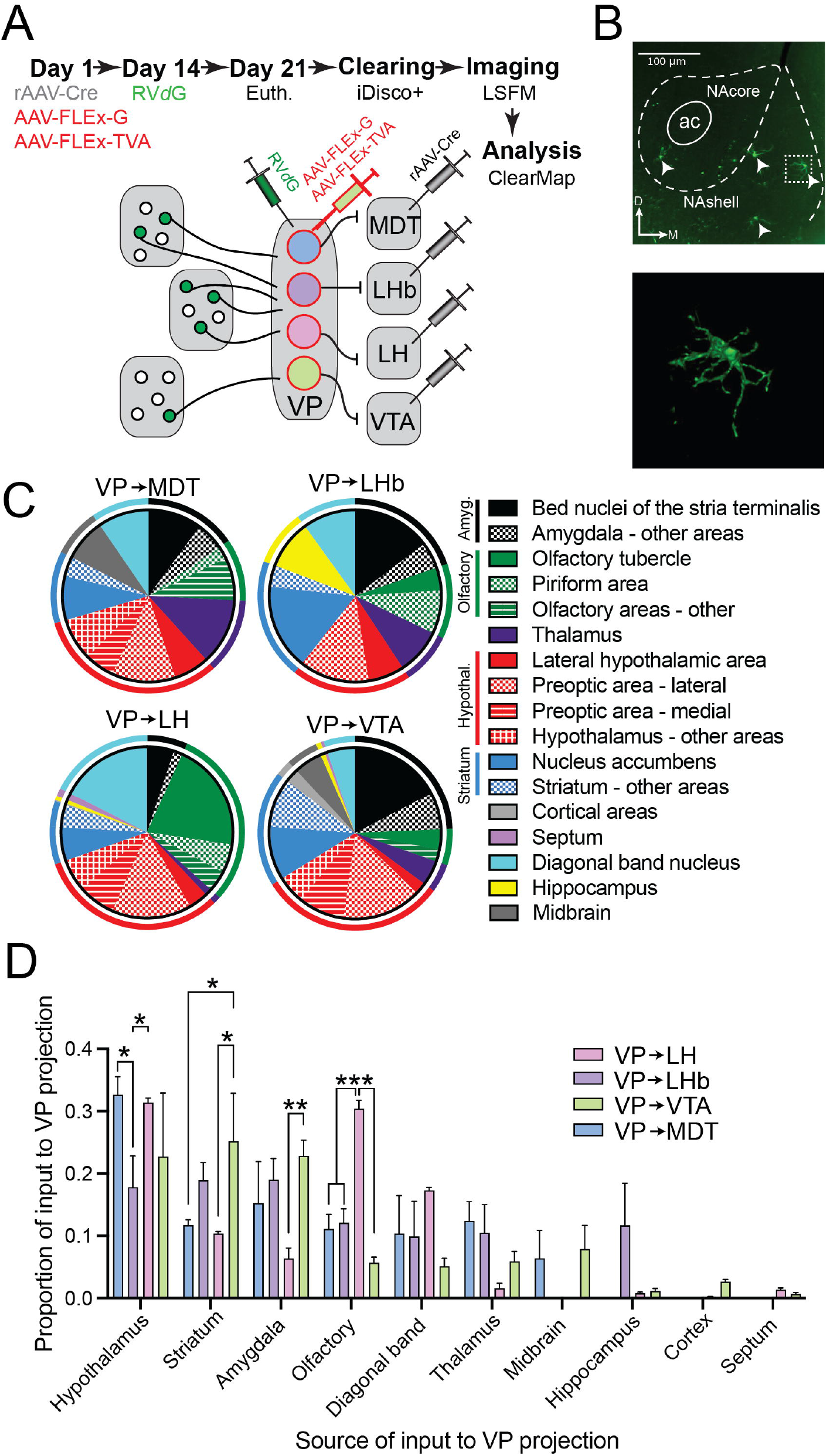
Distinct monosynaptic input patterns between different ventral pallidal projections. (**A**) To visualize the monosynaptic inputs to each ventral pallidum projection we injected a retrograde virus expressing Cre (rAAV-Cre) into one of the targets of the VP and the rabies helper viruses expressing in a cre-dependent manner the avian tumor receptor A (AAV-FLEx-TVA) and the rabies glycoprotein (AAV-FLEx-G) into the VP. Fourteen days later we injected the pseudotyped rabies virus (RV*d*G) into the VP to label all monosynapic inputs to a certain VP projection. We then cleared the brains with iDisco+ and imaged the cleared brains with light sheet fluorescence microscopy (LSFM). (**B**) Top - LSFM image of labeled neurons in the NAc. Bottom – magnification of the neuron marked by a dotted square in the top image. (**C**) Proportions of the major inputs to the four VP projections. Colored ring surrounding pie charts indicates grouping of subparts of the same brain region into the regions used in panel (D). Although the four VP projections receive input from the same brain regions, the proportions of each input vary between the projections. (**D**) The proportions of specific inputs between the four VP projections are significantly different (Two-way ANOVA, main effect for input source, F_(9,80)_=23.8, p<0.0001; interaction (input source X VP projection) effect, F_(27,80)_=3.42, p<0.0001). The VP_→LH_ projections show several differences from the VP_→VTA_ projections, including stronger olfactory input (Tukey’s multiple comparison test (p<0.0001) but weaker striatal (p=0.016) and amygdalar (p=0.006) inputs. *, p<0.05; **, p<0.01; ***, p<0.001.

Our data show that although all four VP projections receive input from largely the same sources, they differ in the relative contribution of each input (**Fig. 4C,D; Supplementary Table 3**). The main inputs to all VP projections originated in the hypothalamus, striatum, amygdala and olfactory areas, with lesser contribution from the midbrain, hippocampus and cortex.

Some inputs were more pronounced on specific projections (**Fig. 4D**). For example, hypothalamic inputs were less dominant in VP_→LHb_ neurons while olfactory inputs were significantly stronger on VP_→LH_ projections and striatal inputs (mainly from the NAc) were specifically enriched in VP_→VTA_ projections. Interestingly, the VP_→LH_ and VP_→VTA_ projections, that show minimal overlap (**Fig. 1C**), also show various differences in their inputs, the VP_→LH_ projections recivieng more olfactory and less striatal and amygdalar inputs compared to VP_→VTA_ projections.

#### The VP_→MDT_ projection receives less inhibitory input compared to the other VP projections

The projections of the VP may differ not only in their gene expression (**Figs. 2, 3**) or their input patterns (**Fig. 4**) but also in the characteristics of the synaptic inputs they receive. To examine this, we used the whole-cell patch clamp technique and recorded from each of the four VP projections the evoked input from the NAc and the global spontaneous inputs they receive (**Fig. 5**).

**Figure 5.**
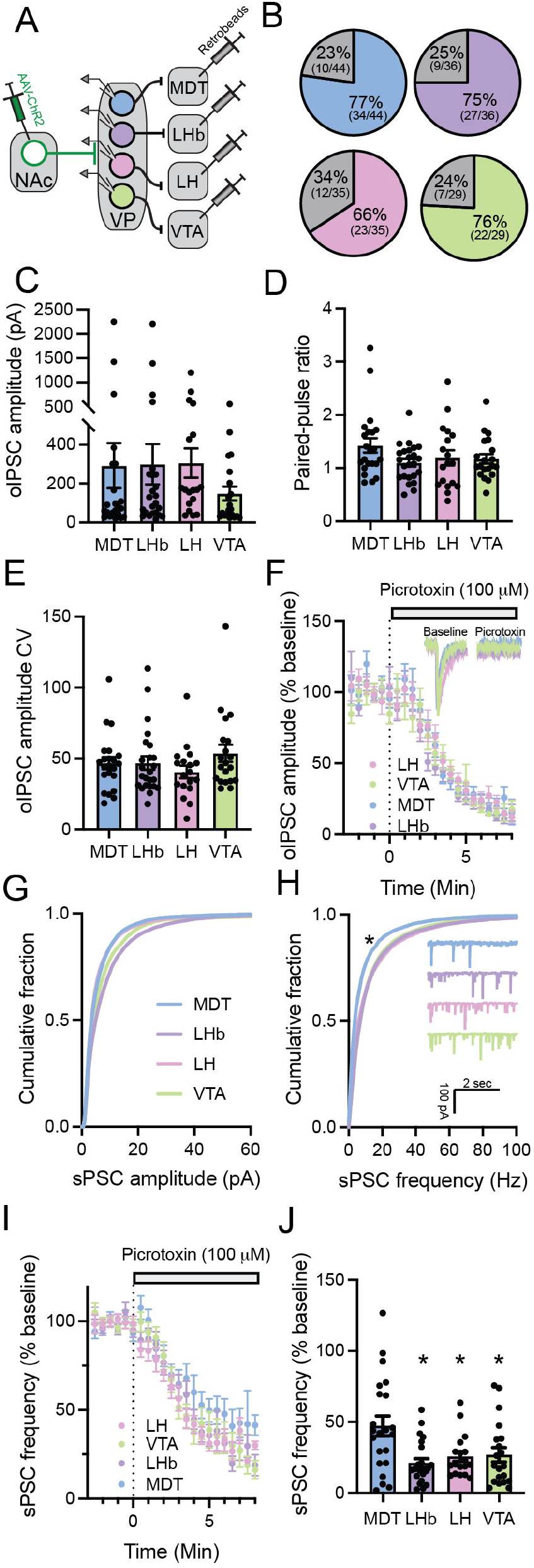
Projection-specific differences in the overall, but not accumbal, inhibitory input to VP neurons. (**A**) Schematic representation of the experimental setup. In each mouse, a retrograde tracer was injected into one of the four targets of the VP examined here and ChR2 was expressed in NAc neurons. Recordings were performed from identified VP projections and NAc input was induced optogenetically. (**B**) The proportion of responding VP neurons (i.e. the number of neurons displaying an oIPSC divided by the total number of neurons examined) was similar between projections and was between 66-77%. (**C-E**) The amplitude (**C**), paired-pulse ratio (**D**) and coefficient of variation (**E**) of the evoked oIPSC did not differ between projections (One-way ANOVA; F_(3,80)_=0.64, p=0.59 for amplitude; F_(3,80)_=1.75, p=0.16 for paired-pulse ratio; F_(3,80)_=1.08, p=0.36 for CV of amplitude). (**F**) Application of 100 μM picrotoxin (GABA-A receptor blocker) on the slice completely blocked the oIPSCs in all VP projections, indicating these are GABA-only inputs. Inset – representative traces. (**G**) Cumulative probability of sPSC amplitudes in the four VP projections. There were no differences between the curves (Kolmogorov-Smirnov tests; p>0.16 for all pairwise comparisons between projections). (**H**) Cumulative probability of the frequency of sPSCs in the four VP projections. The VP_→MDT_ neurons showed significantly lower frequencies compared to the other projections (Kolmogorov-Smirnov tests; d=0.13, p=0.0001 compared to VP_LHb_; d=0.16, p<0.0001 compared to VP_→LH_; d=0.14, p<0.0001 compared to VP_→VTA_). * - p<0.05 for MDT compared with each of the other projections. (**C-D**) Application of 100 μM picrotoxin (GABA-A receptor blocker) on the slice significantly reduced the frequency of spontaneous inputs in all projections (**I**), but the reduction was smaller for the VP_→MDT_ projections (reduction to 47.2±7 % of baseline) compared to all other projections (**J**) [One-way ANOVA main projection effect, F_(3,79)_=4.06, p=0.01; post-hoc Tukey’s multiple comparisons – compared to VP_→LHB_ (21.1±3 % of baseline), p=0.001, compared to VP_→LH_ (25.7±3 % of baseline), p=0.02, compared to VP_→VTA_ (26.9±5 % of baseline), p=0.03]. * - p<0.05 compared to VP_→MDT_ neurons.

Examination of the evoked NAc input to each of the VP projections tested here revealed no difference between the projections. In all projections tested, the proportion of neurons in which we detected evoked NAc input was similar (between 66-77 %, **Fig. 5A-B**) and the average amplitude of the optogenetically-evoked inhibitory postsynaptic current (oIPSC) was similar (**Fig. 5C**). This was also true for the paired pulse ratio (PPR) and coefficient of variation of the evoked currents (two measures reflecting the probability of release^33,34^) (**Fig. 5D-E**). Lastly, in all VP projections application of picrotoxin completely abolished the NAc oIPSC (**Fig. 5F**), indicating a GABAergic synapse.

Comparing the characteristics of the spontaneous postsynaptic currents (sPSCs) between the projections revealed that while they do not differ in the amplitude of the sPSCs (**Fig. 5G**), the VP_→MDT_ projection shows a significantly lower frequency of sPSCs compared to the other projections (**Fig. 5H**). This difference may be due to less GABAergic input because when we washed the slices with picrotoxin (100 μM), a GABA-A receptor antagonist, the reduction in the frequency of sPSCs was significantly smaller in VP_→MDT_ neurons (52.8±33 % reduction) (**Fig. 5I-J**).

### VP_→LH_ neurons are more excitable and VP_→VTA_ neurons are less excitable than the average VP neuron

We next tested various properties that contribute to the excitability of the VP projection neurons (**Fig. 6A**). The resting membrane potential (**Fig. 6B**) did not differ between VP projections and ranged between −45.8 mV and −47.2 mV on average. In contrast, the firing rate at rest (**Fig. 6C-D**) did differ between projections. In particular, the VP_→LH_ neurons had the highest firing rate (10.9±8.1 Hz), which was significantly higher than the firing rate of VP_→MDT_ (6.1±4.8 Hz) and VP_→VTA_ neurons (6.5±5.2 Hz).

**Figure 6.**
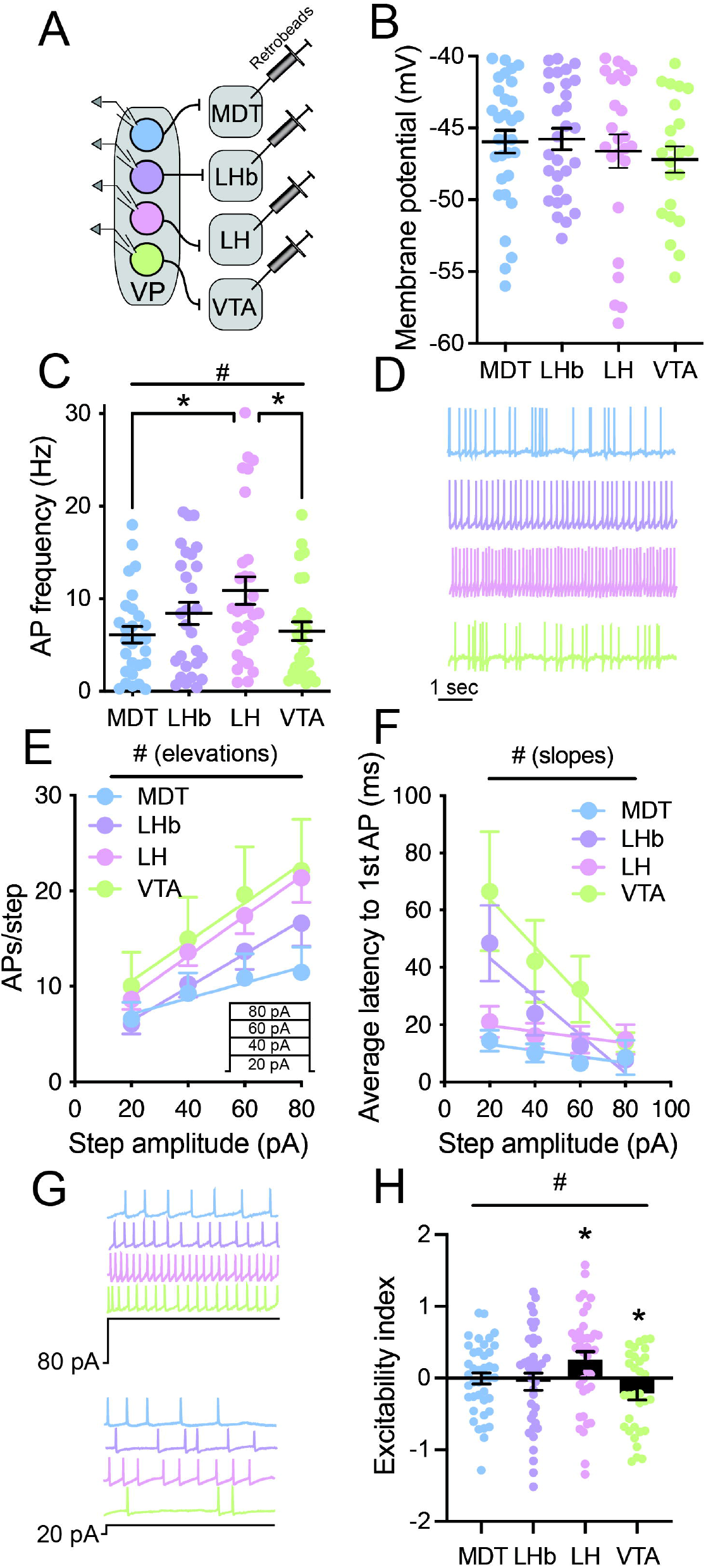
VP_→LH_ neurons are more excitable and VP_→VTA_ neurons are less excitable than the average VP neuron. (**A**) Experimental setup. VP projections were labeled by injecting a retrograde tracer in the target region and recorded from using whole-cell patch clamp electrophysiology. (**B**) Resting membrane potential was not different between the four VP projections we examined (One-way ANOVA, F_(3,103)_=0.49, p=0.69; VP_→MDT_=−46.0±4.4 mV; VP_→LHb_=−45.8±4.0 mV; VP_→LH_=−46.6±5.8 mV; VP_→VTA_=−47.2±4.3 mV). (**C**) Action potential firing frequency at rest differed between the different VP projections (One-way ANOVA, F_(3,103)_=3.47, p=0.02). Post-hoc analyses show that VP_→LH_ neurons fire in higher frequencies than VP_→MDT_ (Post-hoc Holm-Sidak’s multiple comparisons test, p=0.03) and VP_→VTA_ (Post-hoc Holm-Sidak’s multiple comparisons test, p=0.04) neurons. (**D**) Representative traces. (**E-F**) When applying a series of increasing depolarizing steps (inset, 20, 40, 60 and 80 pA) we found a main projection effect both in the number of action potentials per step (Analysis of covariance (ANCOVA), F_(3,260)_=1.06, p=0.37 for slope comparison; F_(3,260)_= 6.1, p=0.0005 for elevation comparison) (**E**) and in the minimal latency to fire the first action potential (ANCOVA, F_(3,260)_=4.65, p=0.003 for slope comparison) (**F**). Note that VP_→LH_ neurons had among the highest number of action potentials and shortest latency to fire, both of which indicate high excitability. (**G**) Representative traces. (**H**) An “Excitability index” was calculated by first calculating for each cell a z-score in each of the parameters recorded in panels B,C,E,F and then calculating the average z-score for each cell. There was a main projection effect for the z-scores (One-way ANOVA, F_(3,146)_=3.21, p=0.03) and post-hoc analyses (Tukey’s multiple comparisons test) revealed that the VP_→LH_ and the VP_→VTA_ projections were significantly different in their excitability (p=0.02). In addition, the excitability index of VP_→LH_ neurons was significantly higher than zero (One sample t-test, t_(36)_=2.18, p=0.035) while that of VP_→VTA_ neurons was significantly lower than zero (One sample t-test, t_(31)_=2.15, p=0.039). Thus, VP_→LH_ neurons are more excitable and VP_→VTA_ neurons are less excitable than the average VP neuron. * - p<0.05 compared to zero.

When we applied a series of depolarizing current steps to the VP projections (**Fig. 6E-G**), we found that the projections differed both in the number of action potentials they generated per step and in the latency to the first action potential. Note that the VP_→LH_ projections, who had the highest firing rates at rest (**Fig. 6C**), also showed strong firing and quick response when depolarized.

To evaluate the average excitability of the projections we generated an “excitability index”. In each of the measured parameters we pooled all cells together and applied z-scores to all cells. Thus, each neuron had four z-scores (one from each experiment) and the excitability index of a neuron is the average of its four z-scores. Comparison of the excitability indexes between VP projections (**Fig. 6H**) demonstrated that it differed between projections and that the VP_→LH_ and VP_→VTA_ projections had significantly different excitability indexes, 0.25 and −0.21, respectively. Moreover, when comparing the excitability indexes to zero, which reflects the average excitability of a VP neuron, the average excitability index of the VP_→LH_ neurons was significantly higher than zero while that of VP_→VTA_ neurons was significantly lower than zero. Thus, our data suggest that VP_→LH_ neurons are a subtype of VP neurons that is more excitable than the average VP neuron while VP_→VTA_ neurons are a differen subtype of VP neurons that is less excitable than the average VP neuron. This, together with our anatomical findings (**Figs. 1, 4**), strengthens the hypothesis that VP_→LH_ and VP_→VTA_ neurons are largely different subpopulations.

#### Genes critical for ion transport enriched within VP_→LH_ and VP_→VTA_ neurons

To identify potential molecules underlying neuronal responses seen between VP_→VTA_ and VP_→LH_ neurons, we compared genes enriched within both VP projection neuron types. We found 156 genes enriched within both VP_→LH_ and VP_→VTA_ neurons (**Fig. 7A**). GO Term analysis illustrated that the 156 genes belonged to processes related to neuron development, synaptic transmission, GABA signaling pathways, chromatin assembly and ion transport (**Fig. 7B**). This is consistent with previous work identifying VP GABAergic projections to the LH and VTA^9,11^. It also suggests that despite electrophysiological differences, there may be shared molecules that regulate presynaptic and synaptic processes between VP_→LH_ and VP_→VTA_ neurons. Given the differences in excitability between these projection neurons, we focused our analysis on genes associated with ion transport, which includes genes encoding subunits of ion channels and transporters. A heat map shows the genes enriched for ion transport within each projection type and their log-fold changes in comparison to global VP neuron gene expression. Among the genes enriched in both VP_→LH_ and VP_→VTA_ neurons, *GABRA5*, the GABA-A receptor alpha 5 subunit, is the highest expressing gene. A predicted transcription analysis on the genes associated with ion transport highlights RFX3 as a top predictor regulator of several shared ion transport genes. This presents a possible target for shared ion genes within VP_→VTA_ and VP_→LH_ neurons.

**Figure 7.**
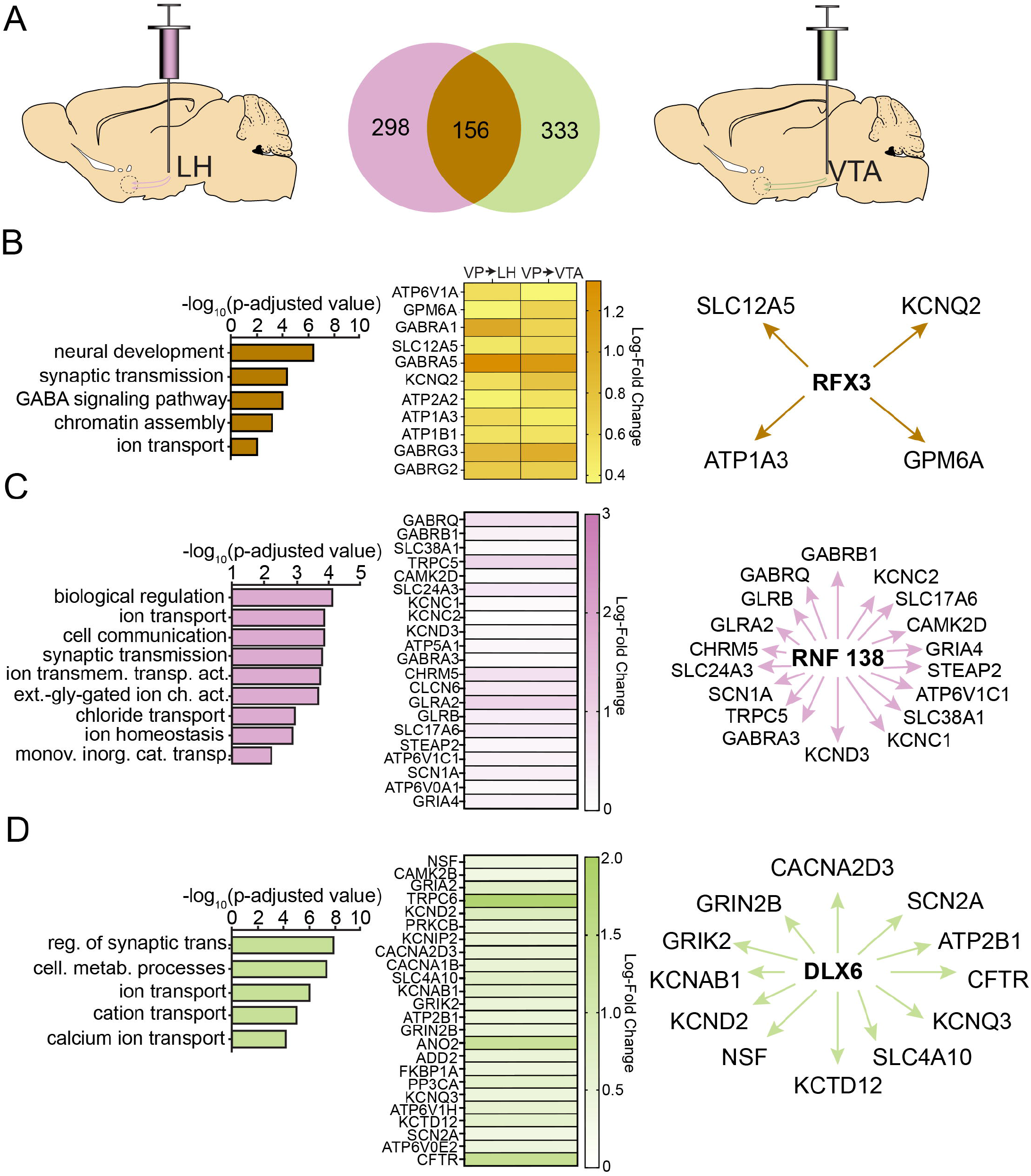
Distinct sets of genes related to ion transport are enriched within VTA-projecting and LH-projecting VP neurons. **(A)** Upregulated genes within VP_→LH_ and VP_→VTA_ projection neurons were compared in a Venn diagram. Gene Ontology analysis identified common synaptic genes (**B**) in VP_→VTA_ and VP_→LH_ projection neurons (log fold changes from each projection type displayed in a heat map) and RFX3 as a predicted upstream regulator of a subset of genes related to ion transport. GO term analysis reveals distinct sets of ion transport-related genes and their listed potential upstream regulators within the VP_→LH_ (**C**) and VP_→VTA_ (**D**) projection neurons. ion transmem. transp. act. – ion transmembrane transporter activity, ext.-gly—gated ion ch. act – extracellular-glycine-gated ion channel activity, monov. Inorg. Cat. Transp. – monovalent inorganic cation transporter, reg. of synaptic trans. – regulation of synaptic transmission, cell. Metab. Processes – cellular metabolic processes.

To identify genes and biological processes distinct to each neuronal type, GO term analysis was performed on the unshared genes within VP_→LH_ (298 genes) and VP_→VTA_ (333 genes) neurons. Interestingly, both sets of genes were associated with processes related to synaptic transmission and ion transport but also show intriguing differences that may contrtibute to the differences in excitability (**Fig. 7C,D**). For example, both neuron types have enriched expression of potassium chanels, despite distinct families expressed within each neuronal type. VP_→LH_ neurons show increased expression of channels from the Kv3 family (*KCNC1, KCNC2*) while VP_→VTA_ neurons show increasd expression of other potassium channel subtypes (*KCND2, KCNAB1*). Since Kv3 channels are known to be important for high-frequency firing^35–37^, the increased expression of *KCNC1, 2* in VP_→LH_ neurons may underlie their higher excitability.

Predicted transcription factor analysis was performed on synaptic genes enriched within VP_→LH_ neurons and VP_→VTA_ neurons. We found that RNF138 was a top predicted regulator for a subset of VP_→LH_ ion transport genes, and DLX4 was a top predicted regulator for a subset of ion transport genes within VP_→VTA_ neurons. Altogether, these data demonstrate that VP_→VTA_ and VP_→LH_ neurons have sets of ion transport genes that are enriched within each projection neuron type and present novel molecular targets to study VP projection-specific neuronal function.

### Opposite roles for VP_→LH_ and VP_→VTA_ projections in cocaine conditioned place preference

Our data so far indicates different VP projections show different characteristics in many aspects. In particular, the VP_→LH_ and VP_→VTA_ projections emerge as largely separate neuronal populations (**Fig. 1**) with different input patterns (**Fig. 4**), different gene expression (**Fig. 7**) and contrasting physiological properties (**Fig. 6**). In light of these differences between the projections, we next tested whether these projections affect differently the motivation to receive drug reward after abstinence in a cocaine conditioned place preference (CPP) procedure.

To inhibit the activity of a specific projection (e.g. VP_→LH_) we injected a retrograde AAV encoding for Cre recombinase (rAAV-Cre) in the target area to express Cre in the specific VP projection neurons and expressed the inhibitory version of the Designer Receptor Exclusively Activated by Designer Drugs (DREADDs) in a cre-dependent manner in the VP (AAV-DIO-hM4Di-GFP) (control mice were injected with a sham virus, AAV-DIO-eYFP) (**Fig. 8A**). We then trained mice to associate one side of the CPP box with cocaine using the cocaine CPP paradigm (**Fig. 8B**, see Methods) and then allowed them to undergo abstinence in their home cages for 14 days. Thirty minutes before the CPP test, we injected all mice with CNO (3 mg/kg, i.p.) and then tested their preference to the cocaine-paired side.

**Figure 8.**
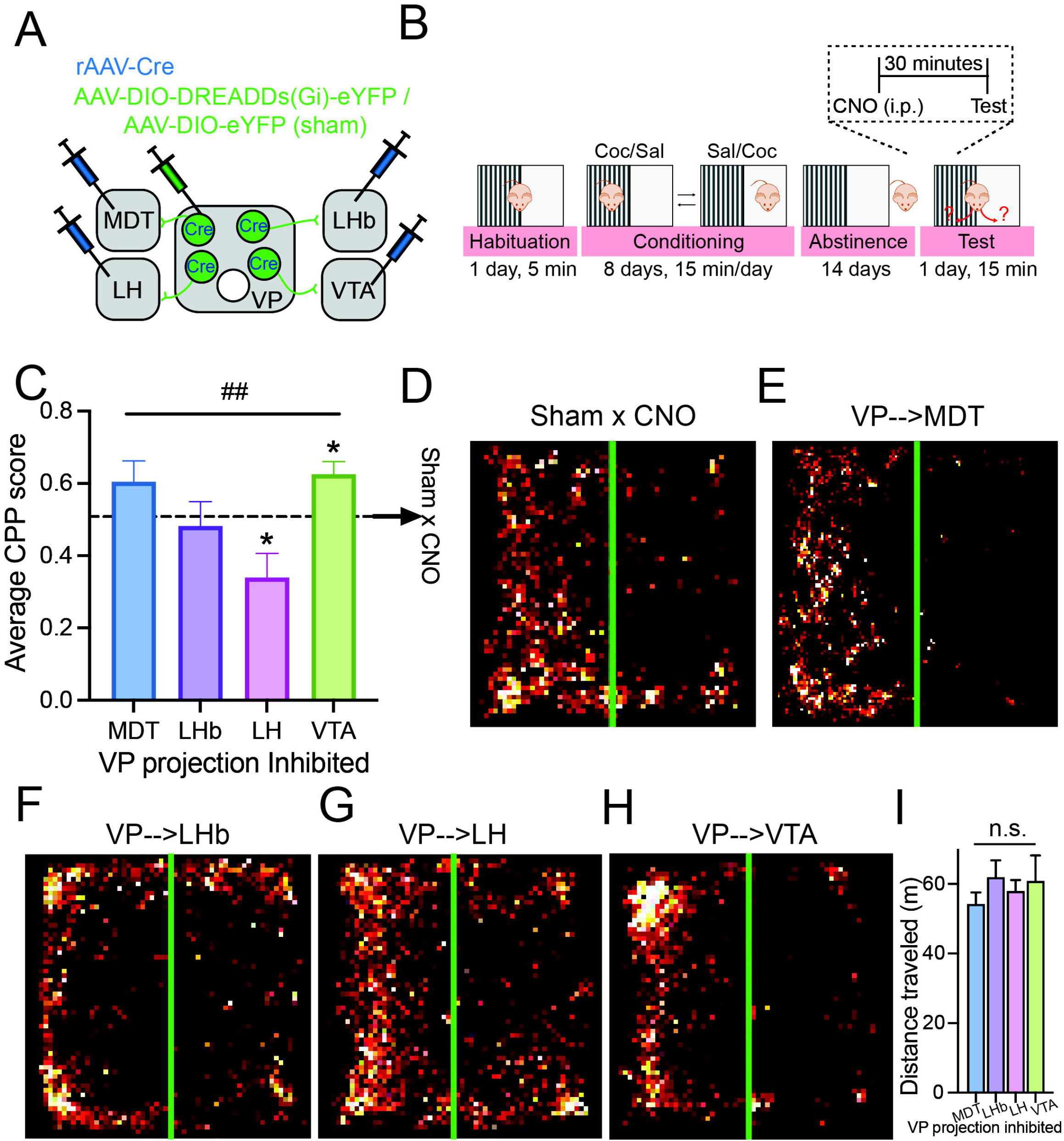
Inhibiting VP_→LH_ neurons diminishes, while inhibiting VP_→VTA_ neurons enhances the preference for cocaine. (**A**) Viral injection strategy. A retrograde AAV expressing Cre (rAAV-Cre) was injected into one of the 4 targets of the VP tested here and an AAV encoding Cre-dependently for the inhibitory DREADD hM4Di (AAV-DIO-hM4Di-eYFP) or a control sham virus (AAV-DIO-eYFP) were injected into the VP. (**B**) Cocaine CPP paradigm. After habituating to the CPP box, mice received daily alternating injections of cocaine (15 mg/kg i.p.) in one side of the box or saline in the other side of the box (4 injections of each). Mice then underwent 14 days of abstinence in their home-cages. On the test day, all mice first received an injection of CNO (3 mg/kg i.p.) 30 minutes prior to the beginning of the test and then were left in the CPP box for 15 minutes, during which their movement was recorded. A CPP score was calculated using the following equation - 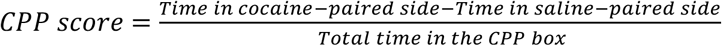. (**C**) Inhibiting different VP projections had different influences on behavior (One-way ANOVA, F_(3,25)_=5.81, p=0.004). While inhibiting VP_→MDT_ or VP_→LHb_ neurons did not affect the CPP score (One-sample t-test comparing to the CPP score of control mice expressing the sham virus and receiving CNO, Sham X CNO, 0.51; t_(6)_=1.71, p=0.14 for VP_→MDT_, t_(5)_=0.38, p=0.72 for VP_→LHb_), inhibiting the VP_→LH_ projection decreased the CPP score (t_(7)_=2.54, p=0.039) and inhibiting the VP_→VTA_ projection increased the CPP score (t_(7)_=3.42, p=0.011). (**D-H**) Heatmaps of the movement of representative mice from all groups. (**I**) Inhibiting the different VP projections did not affect the distance traveled differently between projections.

Our data show that inhibiting the different VP projections had different effects on the preference of the cocaine paired side (**Fig. 8C-H**). In particular, the average CPP score when inhibiting the VP_→LH_ projection (0.34±0.19) was significantly lower than the CPP score when inhibiting the VP_→VTA_ (0.63±0.10) or VP_→MDT_ (0.61±0.15) projections. Comparison of the CPP scores to that of the control group (injected with CNO but expressing the sham virus, average CPP score of 0.51) revealed that inhibiting the VP_→LH_ projection significantly reduced the CPP score while inhibiting the VP_→VTA_ projection significantly increased the CPP score. Inhibiting the VP_→MDT_ or VP_→LHb_ projections did not affect the CPP score. Inhibiting the different projections did not change differently the overall locomotion of the mice (**Fig. 8I**). Thus, our data demonstrates that different VP projections have different roles in renewal of cocaine CPP after abstinence, and highlight the different, and maybe opposite roles of VP_→LH_ and VP_→VTA_ in cocaine preference.

## Discussion

The VP is a structure largely composed of GABAergic neurons and has numerous downstream projection targets. It has been assumed that the cellular composition of the different projections is largely similar, despite the VP’s projection-specific roles in behavior. In this manuscript we provide for the first time a thorough examination of four major outputs of the VP and compare directly their genetic profile, input sources, physiological characteristics and roles in cocaine motivation after withdrawal. We find that the different VP projections show distinct characteristics in all properties examined. Each projection has distinct sets of enriched genes (**Figs. 2–3**), receives different patterns of inputs from various regions of the brain (**Fig. 4**), exhibits different levels of excitability and inhibitory input (**Figs. 5–6**) and has differential roles in regulating cocaine CPP behavior after abstinence (**Fig. 8**). In particular, the VP projections to the LH and VTA stand out as being largely distinct cell populations. They show a minimal level of projection overlap (**Fig. 2**), differences in input sources (**Fig. 4**), express different sets of enriched genes (**Fig. 7**), have different excitability levels (**Fig. 6**) and show opposite effects on cocaine CPP (**Fig. 8**).

### VP heterogeneity

Our data suggest that the four projection neurons of the VP examined here differ from each other in multiple aspects, potentially suggesting some of these projections contain distinct subtypes of VP neurons. This supports previous studies attributing different behavioral roles to distinict VP outputs^5,8,11,29,38,39^. Out of the projections examined here, the VP_→LH_ neurons have the lowest overlap with other VP projection neurons, show enrichment of genes critical for axogenesis and GABAergic receptors and are hyperexcitable. This suggests that VP_→LH_ neurons are a distinct subgroup of neurons that are potentially poised to propagate information quickly from their inputs. In addition, VP_→VTA_ and VP_→LHb_ neurons are enriched for biological processes distinct from other VP projection neurons (VP_→VTA_: Glutamate receptor activity; VP_→LHb_: dopamine binding activity) suggesting these neuronal types may have distinct inputs from other VP projection neurons.

Unlike VP_→LH_ neurons, VP_→MDT_ neurons show an average level of excitability, have the largest proportion of double-labeling with other VP projections and display large overlap of gene expression patterns with other VP projection neuron types. Despite the similarity to other VP projections, VP_→MDT_ neurons are distinct in their low inhibitory input (**Fig. 5**). This relatively weak inhibition of VP_→MDT_ neurons may stem from the small proportion of inputs from the striatum (**Fig. 4**). Thus, a group of neurons may be similar to other neurons in one aspect but different in other aspects, calling for multimodal characterization of neuronal subtypes.

One of the major inputs to the VP is inhibitory input from the NAc. Our data show that about 75% of the neurons in the VP indeed receive synaptic inhibitory input from the NAc (**Fig. 5**), similar to estimations in previous studies^40,41^. Our GO term analysis could reflect this, as all VP projection neurons have genes enriched for GABA receptor activity. Nevertheless, our viral tracing experiments show that more striatal neurons converge on VP_→VTA_ neurons than on VP_→LH_ or VP_→MDT_ neurons (**Fig. 4**). It is possible that exposure to reward may change the innervation pattern of the VP by the NAc, as we and others have previously observed synaptic adaptations in VP neurons after exposure to cocaine or highly-palatable food^24,25,41–43^. Our data overall suggests that VP projection neurons, with the exception of VP_→MDT_ neurons, are largely non-overlapping and composed of distinct genetic and cellular profiles. These findings also highlight the need to further interrogate how the VP encodes distinct sets of information from its various inputs to promote stimuli and input-specific behaviors.

### The difference between VP_→LH_ and VP_→VTA_ projections

Our results highlight the VP_→LH_ and the VP_→VTA_ projections as being particularly distinct from each other. They show only 7% overlap (**Fig. 1**) and a large number of upregulated genes within these populations are non-overlapping (**Fig. 7**). We also detected differences in excitability, with VP_→LH_ neurons being more excitable than VP_→VTA_ neurons (**Fig. 6**), possibly due to specific upregulation of genes encoding for the Kv3 potassium channel family, known to allow high firing frequencies^35–37^ (**Fig. 7**). Consistent with this, chemogenetic inhibition of these populations resulted in opposite behavioral effects; inhibition of VP_→LH_ neurons decreases while inhibition of VP_→VTA_ neurons increases cocaine CPP.

The differences we report on VP_→LH_ and VP_→VTA_ neurons challenge several previous notions of the VP. First, an anatomical study suggests that approximately 90% of VP projections, including those to the VTA, pass through the LH^28^. In fact, the LH is positioned between the VP and the VTA^44^. Our data suggest that the VP_→VTA_ projections do not collateralize to the LH, and are strictly passing through the structure. Our CPP work does demonstrate that the VP exerts projection-specific effects on behavior, as previously seen^5,8,9,45^. However, there is conflicting reports on the effects of inhibiting VP projection neurons on drug seeking. These differences may be due to different cell-type specific manipulations, as Prasad et al. had found that inhibition of VP_→LH_ GABAergic neurons, but not VP_→LH_ parvalbumin-expressing neurons reduces renewal of operant-based alcohol seeking in rats^11^. This may also be due to species-specific effects, as inhibition of VP_→VTA_ neurons has previously been shown to enhance cocaine seeking in mice^5^, but impair cocaine-seeking in rats^46^. Alternatively, it is possible that different circuits and cell-types are affected due to extinction training^11,46^ and forced abstinence^5^. Further investigation into these underlying differences is required to better understand the role of VP in drug-seeking.

The mechanism allowing the VP_→LH_ and the VP_→VTA_ projections to have different behavioral roles in drug reward after abstinence is still not understood. One possibility is potential opposite effects on dopamine release from VTA neurons. VP_→VTA_ neurons target both GABA and dopamine neurons^25,47,48^. It is still not known how activation of VP input to the VTA affects dopamine levels but it is plausible that it enhances dopamine release as it increases the firing rate of putative dopamine neurons^46^. LH GABAergic neurons that project to the VTA increase firing of VTA dopamine neurons^49^. These neurons are also a main target of VP_→LH_ projections^50^. Thus, activation of VP_→LH_ neurons, which are mostly GABAergic^1^, may result in a net decrease in dopamine release. Therefore, VP_→VTA_ and VP_→LH_ are expected to have opposite effects on dopamine release from the VTA. This may underlie their opposite effect on cocaine CPP.

Altogether, our multimodal interrogartion of the VP demonstrates that VP projection neurons have relatively distinct pathways, cellular properties and molecular compositions. These baseline differences may reflect their projection-specific roles of cocaine-related behaviors. Our behavioral findings illustrate how VP projection neurons have differential roles of in cocaine reward, which is consistent with previous work in the field. This work emphasizes the importance in further investigating and characterizing the VP projections and cell types in addiction processes.

## Acknowledgements

This study was supported by the U.S.-Israel Binational Science Foundation (grants 2015239 and 2017252 to YMK and MKL) and the Israeli Science Foundation (Grant No. 1381/15 to YMK).

## Notes

### Competing Interest Statement

The authors have declared no competing interest.

### Summary of Updates

New data from rabies tracing study (Figure 4), reorganization of figures.

