## Supplementary Figure 1 for "Multimodal interrogation of ventral pallidum projections reveals projection-specific signatures and opposite roles in cocaine withdrawal"

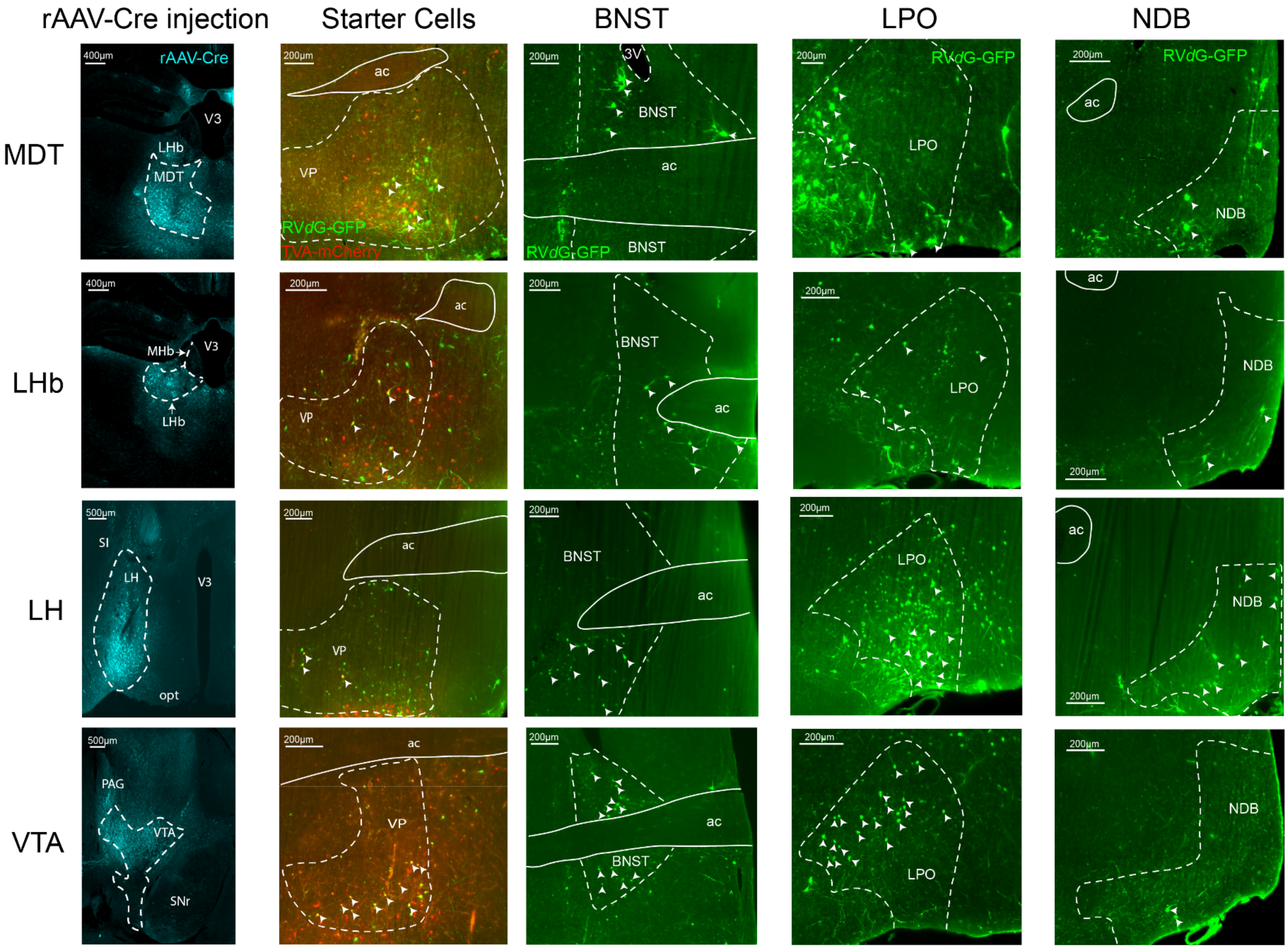
rAAV-Cre injection
Starter Cells
BNST
LPO
NDB
MDT
LHb
LH
VTA
rAAV-Cre
RVdG-GFP
TVA-mCherry
400μm
500μm
200μm
V3
LHb
MDT
MHb
SI
LH
opt
PAG
VTA
SNr
ac
VP
3V
BNST
LPO
NDB

**Supplementary Figure 1 – Injection sites and labeling of inputs to VP projections using the TRIO method.** To label the monosynaptic inputs to each of the four VP projections we tested here, we injected a retrograde AAV expressing Cre into the MDT, LHb, LH or VTA (left column; here we depict injection of rAAV-GFP pseudocolored to cyan to demonstrate injection accuracy) and viruses expressing in a cre-dependent manner the TVA receptor linked to mCherry and the rabies glycoprotein RG into the VP. We also injected the VP with the modified rabies virus (RV*d*G) expressing GFP. Neurons in the VP that were infected with both the TVA receptor (red) and the RV*d*G (green) were considered as starter cells (yellow) (second column from left, arrows point to starter cells). Rabies-labeled neurons in the BNST, LPO and NDB are shown in the 3 rightmost columns.
