## Supplementary Table 3 for "Multimodal interrogation of ventral pallidum projections reveals projection-specific signatures and opposite roles in cocaine withdrawal"

**Supplementary Table 3 - Number of rabies-labeled neurons in main inputs to each VP projection**

| | VP$_{\rightarrow MDT}$ | | | VP$_{\rightarrow LHb}$ | | | VP$_{\rightarrow LH}$ | | | VP$_{\rightarrow VTA}$ | | |
|---|---|---|---|---|---|---|---|---|---|---|---|---|
| Bed nuclei of the stria terminalis | 18 | 0 | 20 | 13 | 38 | 14 | 31 | 65 | 11 | 34 | 87 | 19 |
| Amygdala - other areas | 11 | 4 | 5 | 5 | 12 | 5 | 22 | 7 | 6 | 10 | 40 | 6 |
| Olfactory tubercle | 0 | 0 | 0 | 10 | 5 | 4 | 198 | 154 | 83 | 5 | 18 | 8 |
| Piriform area | 4 | 2 | 3 | 13 | 19 | 4 | 52 | 40 | 22 | 0 | 0 | 0 |
| Olfactory areas - other | 8 | 13 | 9 | 0 | 0 | 0 | 27 | 20 | 37 | 2 | 15 | 1 |
| Thalamus | 19 | 7 | 21 | 16 | 5 | 17 | 6 | 7 | 14 | 11 | 15 | 12 |
| Lateral hypothalamic area | 4 | 9 | 12 | 10 | 18 | 2 | 35 | 27 | 15 | 3 | 14 | 1 |
| Preoptic area - lateral | 10 | 15 | 20 | 10 | 39 | 9 | 144 | 112 | 61 | 11 | 99 | 10 |
| Preoptic area - medial | 8 | 7 | 6 | 0 | 0 | 0 | 66 | 51 | 29 | 4 | 64 | 2 |
| Hypothalamus - other areas | 9 | 7 | 14 | 0 | 0 | 0 | 58 | 47 | 25 | 5 | 34 | 4 |
| Nucleus accumbens | 10 | 8 | 13 | 17 | 44 | 10 | 60 | 47 | 25 | 36 | 30 | 15 |
| Striatum - other areas | 3 | 2 | 9 | 11 | 4 | 3 | 41 | 32 | 17 | 29 | 21 | 25 |
| Cortical areas | 0 | 0 | 0 | 0 | 0 | 0 | 3 | 1 | 1 | 6 | 12 | 3 |
| Midbrain | 1 | 3 | 26 | 0 | 0 | 0 | 0 | 0 | 0 | 12 | 10 | 21 |
| Hippocampus | 0 | 0 | 0 | 7 | 10 | 24 | 7 | 4 | 5 | 3 | 2 | 2 |
| Septum | 0 | 0 | 0 | 0 | 0 | 0 | 11 | 9 | 8 | 2 | 1 | 1 |
| Diagonal band nucleus | 2 | 22 | 12 | 30 | 11 | 3 | 168 | 131 | 71 | 5 | 27 | 10 |

- Each column represents a different hemisphere. N=3 hemispheres for each VP projection.
